# Disruption of mitochondrial integrity impairs early lineage specification and embryonic development in the novel conditional MitoAtDarT mouse model

**DOI:** 10.1101/2025.07.14.664670

**Authors:** Mona Ahmed, Niyati Gadepalli, Gayathri PS, Maroof Hashmi, Anjana Badrinarayanan, Raj K Ladher

**Author notes:** Author for Correspondence: Raj K. Ladher. MA - EMBL Australia Node in Single Molecule Science, Department of Molecular Medicine, School of Biomedical Sciences, University of New South Wales, Sydney, NSW 2052, Australia. GPS - Indian Institute of Technology Hyderabad, Kandi, Telangana 502284, India.

## Abstract

Mitochondria are increasingly recognised as regulators of developmental state, but how mitochondrial dysfunction influences lineage transitions during mammalian embryogenesis remains poorly understood. Here, we generate a conditional mouse model expressing a mitochondrially targeted, attenuated variant of the DNA ADP-ribosyltransferase DarT (mitoAtDarT). Early activation of mitoAtDarT reduced mitochondrial membrane potential and mitochondrial area, altered mitochondrial morphology, and caused embryonic lethality, without detectable increases in nuclear γH2AX. Developmental defects were evident before implantation: mitoAtDarT expressing blastocysts contained fewer cells and an increased proportion of CDX2/SOX2 double-positive cells, consistent with delayed or incomplete resolution of trophectoderm and inner-cell-mass identities. Following implantation, embryos showed progressive growth retardation and reduced proliferation without widespread apoptosis. By E7.5, Brachyury-positive posterior mesoderm was retained and expanded despite failure of allantoic morphogenesis, while broad SOX2- and SOX17-positive compartments remained recognisable. These defects were accompanied by increased PKM2 expression. Together, our findings suggest that mitochondrial dysfunction can compromise the resolution and progression of developmental states. The conditional mitoAtDarT model provides a means to determine whether mitochondrial state represents a general requirement for lineage transitions or whether particular developmental programmes are selectively vulnerable.

## INTRODUCTION

Embryonic development can be considered a series of progressive cellular state transitions. Embryonic cells traverse a series of transient developmental programmes, successively handing off one cellular identity while acquiring the next to generate the different tissues of the organism (Moris et al., 2016). These transitions are coordinated in both space and time, with developmental programmes not only initiated but also terminated appropriately to permit progression toward increasingly specialised cell fates (Miroshnikova et al., 2023). Failure to exit these transient states can be as detrimental as failure to establish them, resulting in inappropriate maintenance of progenitor identity, defective morphogenesis and embryonic arrest. Entry and exit from these states require inputs from signalling pathways, transcriptional networks and chromatin regulation (Gokbuget and Blelloch, 2019). Growing evidence suggests that cellular metabolism is an additional component of this regulatory environment (Cao et al., 2024; Miyazawa and Aulehla, 2018), but how metabolic state integrates with these developmental inputs remains less well understood.

Beyond their role in ATP production through oxidative phosphorylation (OXPHOS), mitochondria can integrate bioenergetics with calcium homeostasis, reactive oxygen species (ROS) signalling, apoptosis and epigenetic regulation (Chandel, 2014; Rizzuto et al., 2012). Through the production of metabolites such as α-ketoglutarate, acetyl-CoA and NAD⁺, mitochondria directly regulate chromatin remodelling, DNA methylation and transcription factor activity, providing a mechanism by which metabolic state can influence developmental competence (Harvey et al., 2019). Through these activities, mitochondria can support the energetic demands of development and influence cells as they transition between successive states (Chakrabarty and Chandel, 2021). This raises the possibility that mitochondrial function contributes directly to developmental progression rather than simply sustaining it.

Early mammalian embryogenesis offers a useful system for examining this question, as the embryo undergoes a succession of rapid, well-defined developmental transitions. Following fertilisation, cleavage divisions generate the blastocyst, where the first lineage decision separates the trophectoderm (TE), which forms the placenta, from the inner cell mass (ICM), which gives rise to the embryo proper and primitive endoderm (Rossant and Tam, 2009). Implantation around embryonic day (E)4.5 initiates reorganisation of the embryo into the egg cylinder before gastrulation establishes the three germ layers through coordinated Wnt, Nodal, BMP and FGF signalling (Morgani and Hadjantonakis, 2020; Tam and Loebel, 2007). Cells emerging from the primitive streak subsequently migrate, differentiate and contribute to both embryonic and extraembryonic tissues, including the allantois (Beddington and Robertson, 1999; Downs et al., 2009). Throughout these events, cells transition successive developmental states while integrating extrinsic signalling with intrinsic metabolic programmes (Cao et al., 2024).

Consistent with these developmental transitions, mitochondrial architecture and metabolism are extensively remodelled during early embryogenesis (Sharpley et al., 2021; Stern et al., 1971; Zhao et al., 2021). Cleavage-stage embryos contain relatively immature mitochondria with sparse cristae, which mature as development proceeds (Stern et al., 1971). Metabolic pathway utilisation also changes substantially during preimplantation development, with increasing glucose utilisation and mitochondrial activity as embryos approach the blastocyst stage (Houghton et al., 1996; Leese and Barton, 1984; Zhao et al., 2021). By the blastocyst stage, trophectoderm cells develop elongated mitochondria with elaborate cristae and increased oxidative metabolism, whereas mitochondria within the ICM remain comparatively immature, consistent with the distinct metabolic requirements of pluripotent and differentiated lineages (Houghton et al., 1996). More recent studies have demonstrated that these metabolic changes are not simply a consequence of differentiation. In the gastrulating mouse embryo, spatially regulated glucose metabolism controls epiblast fate acquisition and mesoderm migration through interactions with developmental signalling (Cao et al., 2024), while stem-cell-derived embryo models show that glycolytic activity regulates Wnt and Nodal signalling and that the balance between glycolysis and oxidative phosphorylation predicts subsequent developmental outcome (Stapornwongkul et al., 2025; Villaronga-Luque et al., 2025).

Mitochondrial function is dynamically regulated during development through changes in organelle morphology and dynamics, respiratory activity, biogenesis and mitochondrial genome maintenance, allowing mitochondrial output to adapt to changing cellular requirements (Facucho-Oliveira and St John, 2009; Wanet et al., 2015). Perturbation of mitochondrial dynamics can itself alter differentiation, indicating that this remodelling is not necessarily a passive consequence of changing cell state (Kasahara et al., 2013). However, despite increasing appreciation that metabolism contributes to embryonic patterning, it remains unclear to what extent mitochondrial function accommodates the changing demands of development or actively enables cells to progress through successive developmental states. Addressing this question has been challenging because severe disruption of mitochondrial function frequently compromises proliferation, viability, and embryonic survival (Hance et al., 2005; Larsson et al., 1998), making it difficult to distinguish specific effects on developmental progression from the broader consequences of mitochondrial dysfunction.

To investigate the developmental contribution of mitochondrial function, we generated a conditional knock-in mouse model expressing a mitochondrially targeted, attenuated variant of the bacterial DNA ADP-ribosyltransferase DarT (mitoAtDarT), providing a genetically controlled means of inducing mitochondrial DNA damage (Dua et al., 2022; Lawaree et al., 2020). Rather than directly perturbing developmental signalling pathways, this approach allows mitochondrial integrity to be compromised during embryogenesis. We show that mitochondrial dysfunction disrupts resolution of the first lineage decision at the blastocyst stage and subsequently perturbs embryonic growth and gastrulation-associated morphogenesis. These defects are accompanied by altered mesodermal patterning, failure of allantois formation, and sustained Brachyury expression in the caudal mesoderm, consistent with impaired developmental progression. Together, our findings support a model in which mitochondrial function contributes not only to the metabolic demands of embryogenesis but also to the orderly progression of cells through successive developmental states.

## MATERIALS AND METHODS

### Mice

Animal protocols were approved by the NCBS Institutional Animal Care and Use Committee. For whole embyo expression, R26R-mitoAtDarT females were crossed with the Pgk1-Cre (Tg (Pgk1-cre^1Lni/Ncbs^ strain (JAX#020811) (Lallemand et al., 1998).

#### Generation of R26R-mitoAtDarT mice

R26R-mitoAtDarT (B6N; DBA2-Rosa26em1^LSL-mitoAtDarT-FLAG-IRESeGFPRL/Ncbs^) was generated using the CRISPR/Cas9 technique. Briefly, mammalian codon-optimized Su9-AtDarT (E. coli DarT G49D-3x FLAG-tag) sequence was synthesized by GeneArt and inserted into the PR26-CAG GFP vector (a kind gift from Ralf Kuehn - Addgene plasmid # 74286) (Chu et al., 2016).

#### Microinjection and embryo transfer

For microinjection, single-cell embryos were collected from matings of DBA2 males with super-ovulated C57BL/6 N females. Microinjection was performed using standard procedures. The injection solution consisted of 25 ng/µl Cas9 protein, 12.5 ng/µl sgRNA, and 20 ng/µl of the donor plasmid (PR26-mitoAtDarT-EGFP) in the microinjection buffer.

Injected embryos were cultured in KSOM media. At the two-cell stage, embryos were transferred to the oviduct of pseudopregnant females. The efficiency of the sgRNA and potential off-target effects were assessed as previously described (Chu et al., 2016).

Genotyping primers are the following

AtDarTF: ATACGAAGTTATCGGCGCGCCATGGCCT;

AtDarTR: CAGGTTGGACACCAGGATCACGAT;

NeoF: GATCGACAAGACCGGCTTCCA;

NeoR: GTTGTCACTGAAGCGGGAAGGGACT.

### Blastocyst collection and staining

Three- to four-week-old female mice were superovulated by intraperitoneal injection of 5 IU pregnant mare serum gonadotropin (PMSG) at approximately 2:00 PM, followed 46–48 hours later by 5 IU human chorionic gonadotropin (hCG). At embryonic day 3.5 (E3.5), around noon, females were euthanized as described previously. Uterine horns were dissected and placed in M2 medium (Sigma). Blastocysts were flushed from the uterine horns using a 26.5-gauge needle attached to a syringe filled with M2 medium. Embryos were collected by mouth pipetting and either maintained in KSOM medium for live staining and imaging or fixed in 4% paraformaldehyde (PFA) for 15 minutes. For whole mount immunostaining, blastocysts were permeabilised, blocked, before incubation with primary antibodies against CDX2 (Biogenix-MU392A) and SOX2 (Invitrogen-14-9811-82) (1:200 dilution in 10% goat serum, 1% BSA in PBST with 0.3% Triton, blocking buffer) overnight at 4°C. Data is from 3 biological replicates: 17 control blastocyst and 15 R26-mitoAtDarT.

### Cryosectioning

Uterine segments containing the embryos were fixed in 4% paraformaldehyde (PFA) for 30 min at room temperature with gentle agitation. Samples were then washed in phosphate-buffered saline (PBS), sequentially equilibrated in 10%, 20%, and 30% sucrose, and embedded in OCT medium (Sakura Chemicals). Embryos were sectioned sagittally at 20 µm using a cryostat (Thermo Scientific HM525NX), and sections were collected onto charged slides. Prior to immunostaining, slides were dried at 37°C for 4–5 h.

### Immunohistochemistry (IHC)

Cryosections were washed with 0.3% PBST three times for 20 minutes each to ensure permeabilization and complete removal of the OCT. Tissues were then blocked in 10% heat-inactivated goat serum (Himedia) and 1% BSA (Sigma) in 0.3 % PBST for 1 hour, followed by overnight incubation with primary antibodies: CdX2 (Biogenix-MU392A), SOX2 (Invitrogen-14-9811-82), SOX17 (R&D systems-af1924), Brachyury (Abcam-EPR18113), TOMM20 (Thermofisher-MA532148), FLAG-tag (Sigma-F1804), PKM2 (Cell Signalling Technology –4053S), phospho-Histone H3 (pHH3) (Cell Signalling Technology - 9706S), and Cleaved caspase (Cell Signalling Technology - 9661S) at a 1:200 concentration diluted in the same blocking buffer, overnight at 4°C. After incubation with primary antibodies, embryos were washed three times in PBST, re-blocked for 1 hour, and subsequently incubated with the respective secondary antibodies (1:1000) for 1 hour. Nuclei were counterstained with DAPI for 7–10 minutes, followed by a final wash in PBST.

Cryosections were mounted using Fluoroshield mounting medium. Preimplantation embryos were loaded onto plates with coverslips for confocal imaging.

### Mitochondrial membrane potential (MMP)

MMP was assessed using TMRE staining. Briefly, freshly dissected E7.5 mouse embryos were incubated with 200 nM TMRE in DMEM high glucose, GlutaMAX, media for 30 minutes in the incubator at 37°C with 5% CO2. Following the treatment, the embryos were washed in HBSS and transferred to a 35-mm dish with a coverslip before being imaged using a confocal microscope. Data were collected from three biological replicates: six R26-mitoAtDarT and six littermates control embryos. For comparison between embryos, the mean TMRE intensity was normalized to the highest value.

### EdU Click Reaction

At embryonic day 7.5 (E7.5), pregnant PGK1-Cre homozygous females crossed with R26R-mitoAtDarT heterozygous males were intraperitoneally injected with 2.5 mg per 50 g of body weight EdU (Invitrogen C10337) four hours before euthanasia. Uterine segments were dissected, fixed, and sectioned as mentioned above. Following permeabilization, the sections were incubated in the Click-iT reaction cocktail (Invitrogen C10337) according to the manufacture for two hours, containing Alexa Fluor® 555 to detect EdU incorporation. The sections were then processed according to the standard immunohistochemistry (IHC) protocol for further analyses. Data were collected from three biological replicates: four R26-mitoAtDarT and five littermates control embryos.

### Western blot

The total proteins were extracted from the remaining mouse embryonic or extraembryonic tissue using RIPA buffer and sonication, quantified by BCA (Bio-Rad) as described, and separated by SDS-PAGE. The blots were probed with anti-FLAG antibody (M2, Sigma) and incubated with secondary antibody anti-mouse HRP (GE Healthcare). The probed blot was visualized using the ECL detection kit (Advansta).

### Mass Spectrometry and Metabolomics

E7.5 mouse embryos were collected from two crosses: PGK1-Cre homozygous females crossed with R26R-mitoAtDarT homozygous males (R26-mitoAtDarT), and B6ND2 females crossed with R26R-mitoAtDarT males (Control). For each condition, we pooled embryos from 3–4 dams and stored them at −80 °C until further processing.

Metabolites were extracted in 60 μL of ice-cold 50% acetonitrile (Fisher Scientific, A955-4) containing 0.1% formic acid and 2 μg/sample D4-alanine (Cambridge Isotope Laboratories, DLM-250-1) as an internal standard. Samples were briefly vortexed and centrifuged at 13,000 rpm for 30 min at 4 °C. The supernatant was collected, and 5 μL was injected for metabolite analysis.

Glycolytic and TCA-cycle metabolites, including acetyl-CoA, were analysed using a Sciex QTRAP 5500 triple-quadrupole mass spectrometer coupled to a Shimadzu UPLC system. Metabolites were separated on an Acquity UPLC BEH HILIC column (2.1 × 100 mm, 1.8 μm; Waters Corp., USA), using an LC protocol described previously (PMID: 41547903; DOI: 10.1038/s41598-026-35476-8). The column temperature was maintained at 35 °C and the autosampler at 5 °C. Mobile phase A consisted of acetonitrile/water (95:5), and mobile phase B of acetonitrile/water (50:50), with both containing 10 mM ammonium acetate (Sigma, 73594-25G-F) and 0.1% formic acid.

The elution gradient was as follows: 99% A from 0–2 min at 0.4 mL/min; 99–45% A from 2–8 min at 0.4 mL/min; 45–1% A from 8–9 min at 0.4 mL/min; 1% A from 9–9.1 min while increasing the flow rate from 0.4 to 0.8 mL/min; 1% A from 9.1–10 min at 0.8 mL/min; 1–99% A from 10–10.1 min at 0.8 mL/min; 99% A from 10.1–11 min at 0.8 mL/min; 99% A from 11–14.1 min while decreasing the flow rate from 0.8 to 0.4 mL/min; and 99% A from 14.1–15 min at 0.4 mL/min.

Mass-spectrometry parameters followed the manufacturer’s recommendations (Phenomenex TN-1241). Multiple-reaction monitoring (MRM) methods were adapted from Walvekar et al. (2018). Peak areas were quantified using AB SCIEX MultiQuant software (version 3.0.1).

Following metabolite extraction, the protein pellet was resuspended in RIPA buffer and protein concentration was determined using a BCA assay. Metabolite measurements were normalised to the protein content of each sample.

### MicroCT scanning and data processing

Tissue processing and microCT imaging were performed using a protocol modified from one previously described (Ermakova et al., 2018). At the indicated embryonic days, the pregnant females were sacrificed, and the uterine segments were collected, fixed, dehydrated, and embedded in paraffin.

The 3D scan of the paraffin-embedded samples was acquired using a micro–CT Bruker Skyscan1272 machine at 40 kV, 150 µA, 4940X3280 pixels, averaging set to 3, with a rotation step of 0.2° for 180°. The image acquisition was performed without filters at 0.7 µm resolution. The acquired 2D images were reconstructed using NRecon Skyscan reconstruction software. Samples were treated with a ring artifact factor of 20 units and smoothing of five units. The stacks were then reoriented and aligned using DataViewer software (Bruker). Each embryo’s coronal view was saved in BMP format and used to draw the region of interest (ROI) around each embryo. The volume and the surface area of the embryos were calculated from the signal-bearing voxels within the drawn ROI using CT analyzer software. The stacks were thresholded using automatic (Quartile) on 2D space with a quantile of 0.3, with a Gaussian blur filter applied. Statistical significance was calculated through the two-tailed Mann–Whitney U test in GraphPad prism.

For movies and 3D rendering, CTvox v 2.5 (Bruker) 3D Visualization Software was used. For E7.5 analysis, data were collected from three biological replicates: twenty-four R26-mitoAtDarT and twenty-seven R26R-mitoAtDarT control embryos. While for e6.5 analysis, data were collected from three biological replicates: ten R26-mitoAtDarT and fourteen R26R-mitoAtDarT control embryos.

### Haematoxylin and Eosin (H&E) staining

Following microCT, same embryos were re-embedded in paraffin, sectioned and mounted on glass slides. Sections were then deparaffinized and rehydrated before stained with H & E using standard procedures.

### Confocal image acquisition and processing

Confocal images were acquired using a point scanning Olympus FV 3000 inverted microscope. For mitoAtDarT colocalization and mitochondrial morphology, images were captured using a 60x oil immersion objective (1.42 numerical aperture) with acquisition setting parameters optimized to meet or below Nyquist sampling criteria. For transcription factors localization analysis, images were acquired with a 20x oil immersion objective (0.85 NA). A few images were acquired just above the Nyquist criteria to cover the entire embryonic region. To achieve an optimum signal, a variable combination of laser power (<10%) was used with PMT voltage (400V – 600V), gain (1), and offset (9). To avoid signal bleed-through between channels, sequential imaging was employed. For live imaging of TMRE-stained embryos, imaging was carried out in a controlled chamber maintained at 37 °C and 5% CO₂.

### Lightsheet imaging

For blastocyst imaging, light-sheet fluorescence microscopy was employed due to its superior tissue penetration and extended working distance. Blastocysts were embedded in glass capillaries and mounted in a sample holder of a Zeiss Light-sheet 7 microscope (Carl Zeiss, Germany). Imaging acquisition utilized Zeiss Zen Black acquisition software: a single plane was imaged using pivot scan mode with 4 to 10% of 25 mW input laser power. For dual-side (left and right) illumination, a pair of 10x air objectives (LSFM 10x/0.2 foc, Carl Zeiss, USA) was employed, while emission was detected orthogonally using a 20x water immersion objective (W Plan Apochromat, 1.0 NA, Carl Zeiss, USA). To improve image quality and reduce shadowing artefacts, online fusion of images from both left and right illuminations was applied during acquisition.

### Image analysis

Acquired images were visualized and analyzed using Imaris 9.3 software (Oxford Instruments) and ImageJ/Fiji (Schindelin et al., 2012). For intensity profile and colocalization analysis, Fiji software with the JaCob plugin was used to calculate the colocalization coefficients. For mitochondria morphology analysis, Fiji software with the Mitochondria Analyzer plugin was utilized. Quantification of antibody-labelled/DAPI-positive nuclei were performed using the Spots module in Imaris.

### Statistical analysis

Data was analyzed and graphs were plotted using GraphPad Prism v9.5.0 software. The statistical significance was calculated using t-tests assuming no Gaussian distribution (Mann-Whitney t-test). *p-*values are plotted in each graph, with *p*<0.05 considered statistically significant.

## RESULTS

### mitoAtDarT: a conditional system for perturbing mitochondrial function through mtDNA damage

To develop a system to spatially and temporally control mitochondrial function perturbation, we adapted an approach based on selective damage to mitochondrial DNA (mtDNA). DarT is a bacterial DNA ADP-ribosyltransferase that modifies single-stranded DNA by mono-ADP-ribosylating specific sequence motifs, thereby interfering with DNA replication (Jankevicius et al., 2016). When targeted to mitochondria in Saccharomyces cerevisiae using the mitochondrial targeting sequence from subunit 9 of the F₁ ATPase (Su9), DarT induces mtDNA damage and mitochondrial dysfunction (Dua et al., 2022).

To adapt this system for use in mammals, we initially attempted to express mitochondrial-targeted DarT from a mammalian expression construct. However, we were unable to generate stable constructs, suggesting toxicity from low-level leaky expression. We therefore used a previously characterised attenuated DarT variant carrying a glycine-to-aspartate substitution at position 49 (G49D) (Lawaree et al., 2020). This variant, hereafter termed AtDarT, retains DNA-targeting activity but has reduced ADP-ribosyltransferase activity, providing a potentially more tractable way to induce mitochondrial DNA damage in mammalian cells.

We generated a conditional knock-in mouse in which mitochondrially targeted AtDarT (mitoAtDarT) was inserted into the Rosa26 locus (Fig. 1A). A guide RNA targeting the first intron of Rosa26 (sgROSA26-1) was used for site-specific integration (Chu et al., 2016); its editing efficiency in our system is shown in Supplementary Fig. 1A, B. The targeting cassette contained a loxP-flanked transcriptional STOP sequence upstream of mammalian codon-optimised AtDarT. To direct AtDarT to the mitochondrial matrix, the Su9 mitochondrial targeting sequence (residues 1–69) was fused to its N-terminus. A 3×FLAG epitope was added at the C-terminus, together with a downstream IRES-eGFP reporter to allow detection of transgene expression (Fig. 1A).

**Figure 1.**
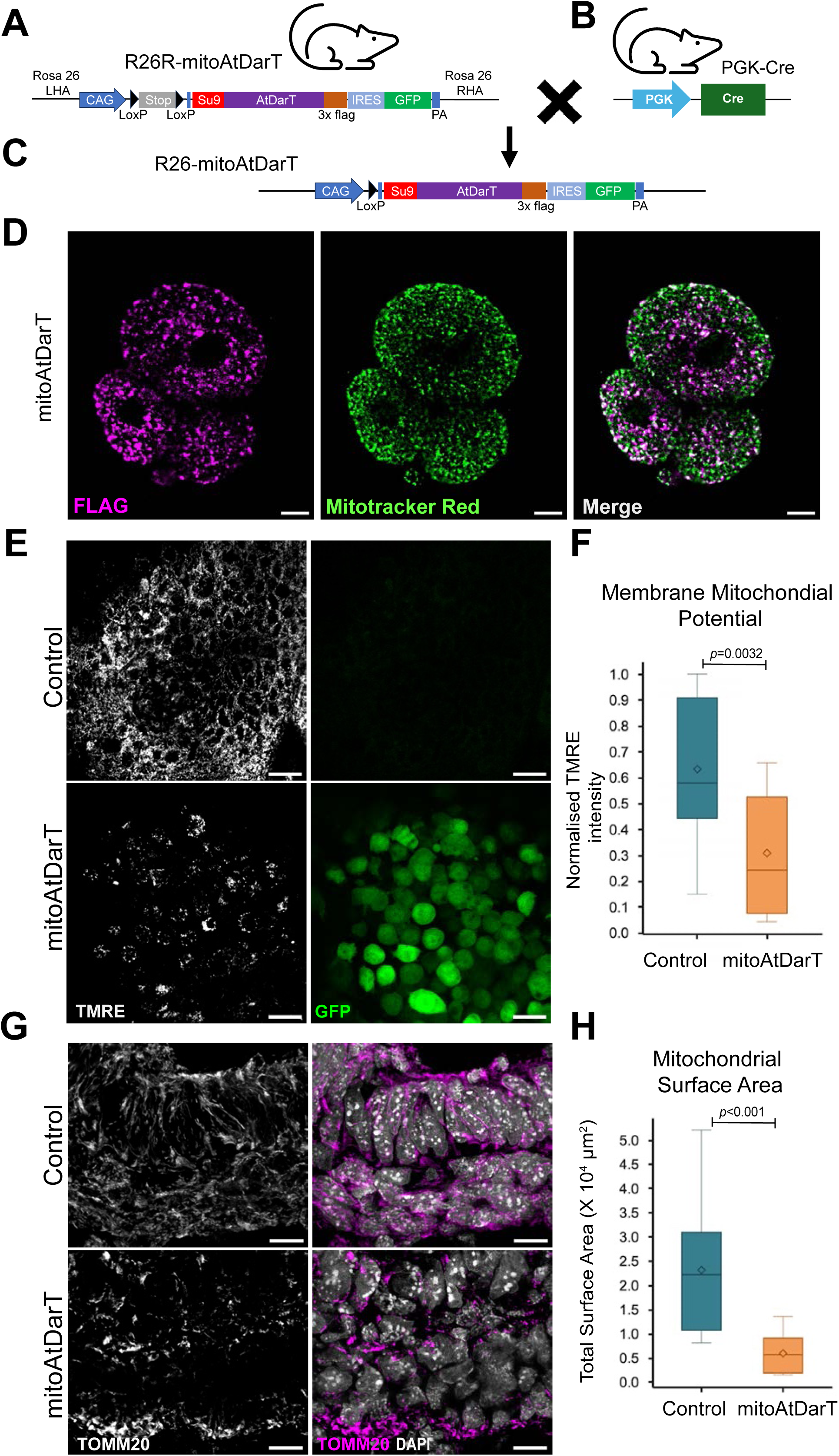
Generation and characterisation of the R26R-mitoAtDarT mouse model. A. Schematic of the R26R-mitoAtDarT targeting construct integrated into the Rosa26 locus. The construct includes a LoxP-STOP-LoxP (LSL) cassette upstream of a mitochondrial targeting sequence (Su9) fused to AtDarT, followed by a FLAG epitope tag, an internal ribosome entry site (IRES), and a GFP reporter. Expression, after LSL excision, is driven by the CAG promoter. B. Schematic of the PGK1-Cre transgenic line, which ubiquitously expresses Cre recombinase under the PGK1 promoter, enabling early and uniform recombination. C. Post-recombination configuration of the R26R-mitoAtDarT allele to generate R26-mitoAtDarT embryos. This results in constitutive expression of mitoAtDarT. D. 4-cell stage embryos showing mitochondrial localization of mitoAtDarT. Mitochondria were stained with MitoTracker Red (pseudo-coloured green) and FLAG-tagged mitoAtDarT, detected with anti-FLAG antibody (magenta). Colocalization analysis revealed a high degree of overlap (Manders’ overlap coefficient = 0.875), confirming proper mitochondrial targeting. Scale bars, 30 μm. E. Mitochondrial membrane potential in whole-mount E7.5 embryos assessed by TMRE staining (GFP, green; TMRE, gray). Scale bars, 20 μm. n = 4 embryos per genotype. F. Quantification of TMRE fluorescence intensity normalized to the highest-intensity control sample. MitoAtDarT-expressing embryos exhibited reduced mitochondrial membrane potential (*p* = 0.0032, Mann–Whitney U t-test). G. Immunostaining of E7.5 embryos for TOMM20, an outer mitochondrial membrane marker (magenta). Nuclei were counterstained with DAPI (gray). Scale bars, 20 μm. n = 4 embryos per genotype. H. Quantification of total mitochondrial area in embryos. MitoAtDarT-expressing embryos displayed a significant reduction in mitochondrial content (*p* < 0.0001, Mann– Whitney U t-test).

Pronuclear microinjection of the targeting construct into fertilised embryos generated three PCR-positive founder animals (#1, #12 and #22), corresponding to a 10% targeting efficiency (Supplementary Fig. 2A, B). Sanger sequencing confirmed the integrity of the AtDarT sequence and surrounding regions (Supplementary Fig. 2C), and two founder lines contained the complete ∼8-kb loxP-STOP-loxP-mitoAtDarT-3×FLAG-IRES-eGFP cassette correctly integrated at the Rosa26 locus. No sequence alterations were detected at the predicted off-target sites examined (Supplementary Fig. 1C–E). In the absence of Cre recombinase, these R26R-mitoAtDarT animals were viable and healthy, with no detectable evidence of transgene expression, establishing a conditional mouse line for subsequent analysis of mitoAtDarT activity.

To activate mitoAtDarT during early embryogenesis, we crossed R26R-mitoAtDarT animals with PGK1-Cre mice. PGK1-Cre is active from the oocyte stage onwards, resulting in early and widespread Cre-mediated recombination (Lallemand et al., 1998). Embryos carrying the recombined allele are referred to hereafter as **R26-mitoAtDarT**. FLAG immunostaining detected mitoAtDarT expression by the 2–4-cell stage, and the protein showed extensive overlap with mitochondrial markers (Manders’ overlap coefficient >0.85; Fig. 1D), confirming targeting to mitochondria.

We next asked whether mitoAtDarT expression was sufficient to compromise mitochondrial function during early development. In freshly isolated E7.5 embryos, we assessed mitochondrial membrane potential using TMRE, a potentiometric dye that accumulates in polarised mitochondria (Scaduto and Grotyohann, 1999). R26-mitoAtDarT embryos showed a marked reduction in TMRE fluorescence compared with controls (Fig. 1E-F), and GFP expression confirmed widespread transgene activation. Mitochondrial abundance and morphology were also altered: TOMM20-positive mitochondrial area was significantly reduced in R26-mitoAtDarT embryos, and mitochondria were smaller, more rounded, and less extensively branched than in controls (Fig. 1G-H; Supplementary Fig. 2D-E). We next asked whether mitoAtDarT expression was associated with detectable nuclear DNA damage using γH2AX, which is recruited to sites of DNA damage. Immunostaining for γH2AX revealed no significant increase in nuclear signal in R26-mitoAtDarT embryos relative to controls, whereas hydroxyurea-treated embryos showed the expected strong γH2AX response (Supplementary Fig. 3). Thus, mitoAtDarT expression did not produce detectable nuclear double-strand DNA damage.

Together, these results suggest the utility of the R26R-mitoAtDarT mouse as a conditional system in which AtDarT can be targeted to mouse mitochondria and produce a mitochondrial phenotype, characterised by reduced membrane potential, reduced mitochondrial content and altered mitochondrial morphology, without detectable nuclear DNA damage. We therefore used this system to examine how disruption of mitochondrial function affects progression through early mammalian development.

### Mitochondrial dysfunction disrupts the resolution of blastocyst lineage identities

As mitoAtDarT expression was detectable from the 2–4-cell stage, we asked whether mitochondrial dysfunction affected development before implantation. We therefore examined R26-mitoAtDarT embryos at embryonic day 3.5 (E3.5), when the first major lineage segregation in the mammalian embryo generates the trophectoderm (TE) and inner cell mass (ICM) (Posfai et al., 2017; Strumpf et al., 2005). Despite forming morphologically recognisable blastocysts, R26-mitoAtDarT embryos contained significantly fewer cells than either R26R-mitoAtDarT or wild-type controls (Fig. 2A-B), indicating that the consequences of mitochondrial dysfunction were already apparent during preimplantation development.

**Figure 2.**
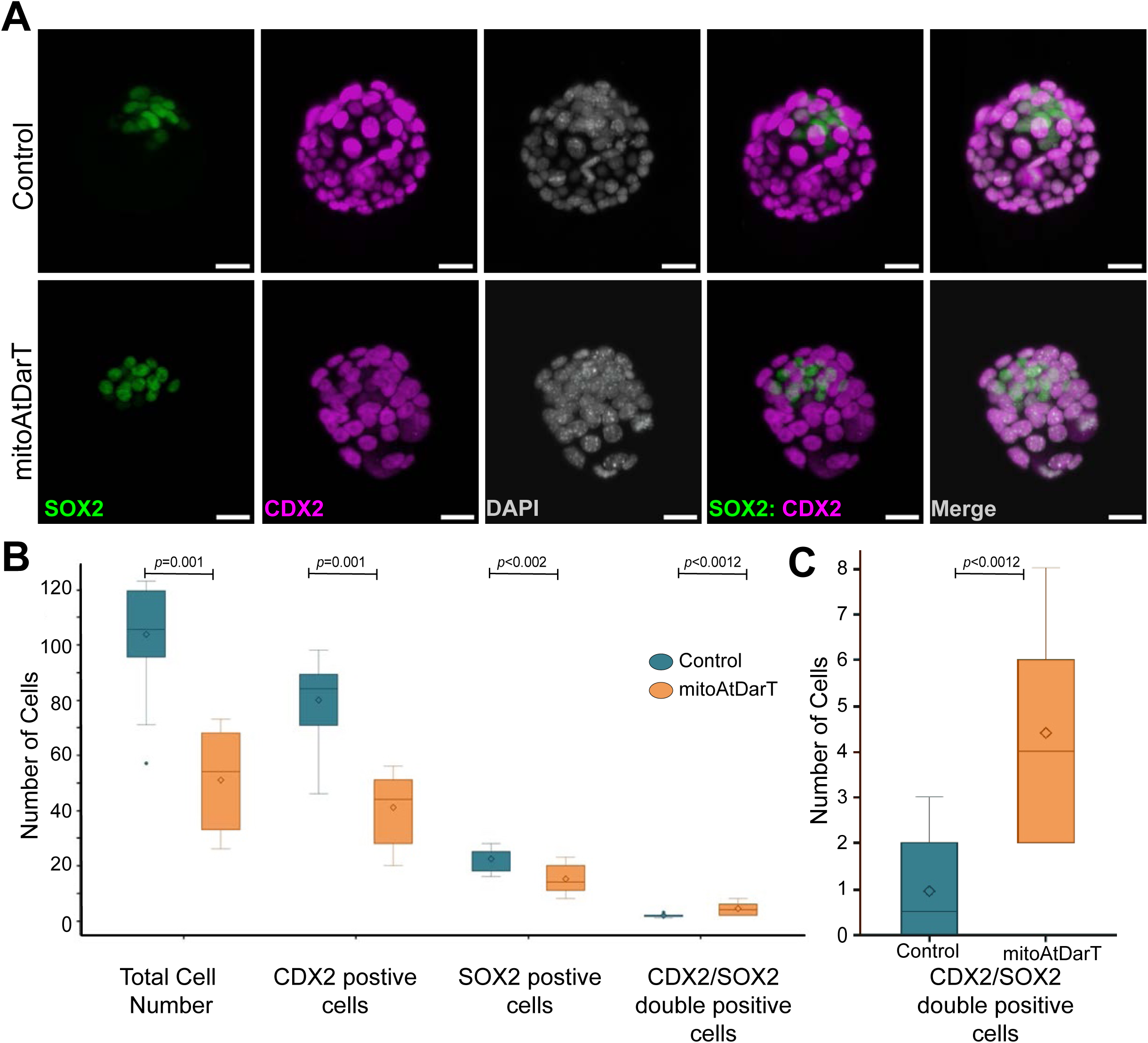
MitoAtDarT-mediated mitochondrial dysfunction disrupts blastocyst formation and lineage segregation. A. E3.5 blastocysts from control (R26R-mitoAtDarT, upper panel) and R26-mitoAtDarT (lower panel) embryos. Immunofluorescence staining shows SOX2 (inner cell mass marker, green), CDX2 (trophectoderm marker, magenta), and nuclei (DAPI, gray). Images were acquired by lightsheet microscopy and analyzed with Imaris software; the Spots module was used for cell quantification. Scale bars, 20 μm. B. Quantification of total cell number (DAPI+) and lineage-specific cells (SOX2+ and CDX2+) per blastocyst. R26-mitoAtDarT embryos show significantly fewer total cells and reduced numbers of SOX2+ and CDX2+ cells. C. Quantification of the number of SOX2/CDX2 double-positive cells per blastocyst in control and R26-mitoAtDarT. Notably, the proportion of cells co-expressing SOX2 and CDX2 is increased in R26-mitoAtDarT embryos. *p* = 0.0012. n=15 R26-mitoAtDarT embryos and 17 R26R-mitoAtDarT (control).

We next asked whether the decrease in cell number was linked to changes in lineage allocation. Blastocysts were stained for CDX2, a marker of the trophectoderm, and SOX2, a marker of the inner cell mass (Posfai et al., 2017; Strumpf et al., 2005). As expected from the overall reduction in cell count, R26-mitoAtDarT blastocysts showed fewer CDX2-positive and SOX2-positive cells compared to controls (Fig. 2B). Nonetheless, the proportions of cells expressing only CDX2 or only SOX2 remained largely consistent, indicating no simple shift toward either TE or ICM lineages. Instead, there was an increase in cells co-expressing CDX2 and SOX2 in R26-mitoAtDarT blastocysts (Fig. 2B, C). These markers normally resolve into their separate domains during blastocyst development (Posfai et al., 2017; Wicklow et al., 2014), but their continued co-expression in mitoAtDarT-expressing embryos suggests delayed or incomplete separation of TE and ICM identities.

Thus, mitoAtDarT-mediated mitochondrial dysfunction does not appear to prevent blastocyst formation or gross allocation of cells to the two major preimplantation compartments. Instead, reduced cell numbers and the persistence of CDX2/SOX2 double-positive cells are consistent with delayed or incomplete resolution of TE and ICM identities. These observations raise the possibility that mitochondrial function contributes to the timely resolution of lineage identity during the first developmental transition of the mammalian embryo.

### mitoAtDarT embryos undergo progressive developmental arrest after implantation

We next examined how the early developmental defects associated with mitoAtDarT expression affected post-implantation development. Crosses between heterozygous R26R-mitoAtDarT males and PGK1-Cre females produced no surviving R26-mitoAtDarT offspring. Mutant embryos could nevertheless be recovered after implantation: at E8.5–E9.5 they were growth-retarded and morphologically abnormal, and by approximately E10 most had undergone resorption. Consistent with this, E11.5–E12.5 implantation sites fell into two distinct classes, comprising normal-sized decidua containing mitoAtDarT-negative embryos and smaller, resorbing decidua, likely associated with mitoAtDarT-positive conceptuses (Supplementary Fig. 4). The frequency of resorbing implantation sites was consistent with the expected Mendelian frequency of mitoAtDarT positive conceptuses.

To define when this phenotype first became apparent, we examined embryos at E6.5 and E7.5 using high-resolution micro-computed tomography (microCT), which allowed embryonic architecture to be visualised within intact uterine segments (Ermakova et al., 2018). At E6.5, R26-mitoAtDarT embryos were already significantly smaller than controls, but retained an elongated egg-cylinder morphology, suggesting that early post-implantation organisation had occurred (Christodoulou et al., 2019)(Supplementary Fig. 5). Quantitative reconstruction confirmed significant reductions in both embryonic volume and surface area at this stage.

The phenotype became substantially more pronounced by E7.5. Control embryos displayed the expected organisation of embryonic and extraembryonic compartments, with clearly identifiable structures including the ectoplacental cone, chorion, amnion and allantois (Fig. 3A, D). In contrast, R26-mitoAtDarT embryos showed considerable morphological heterogeneity, but most were markedly smaller and exhibited disrupted tissue organisation, with poorly defined compartments and collapsed or absent cavities (Fig. 3E, H). Quantitative three-dimensional analysis showed a further reduction in embryonic volume and surface area relative to controls (Fig. 3I, J), while H&E-stained sections confirmed disrupted embryonic architecture.

**Figure 3.**
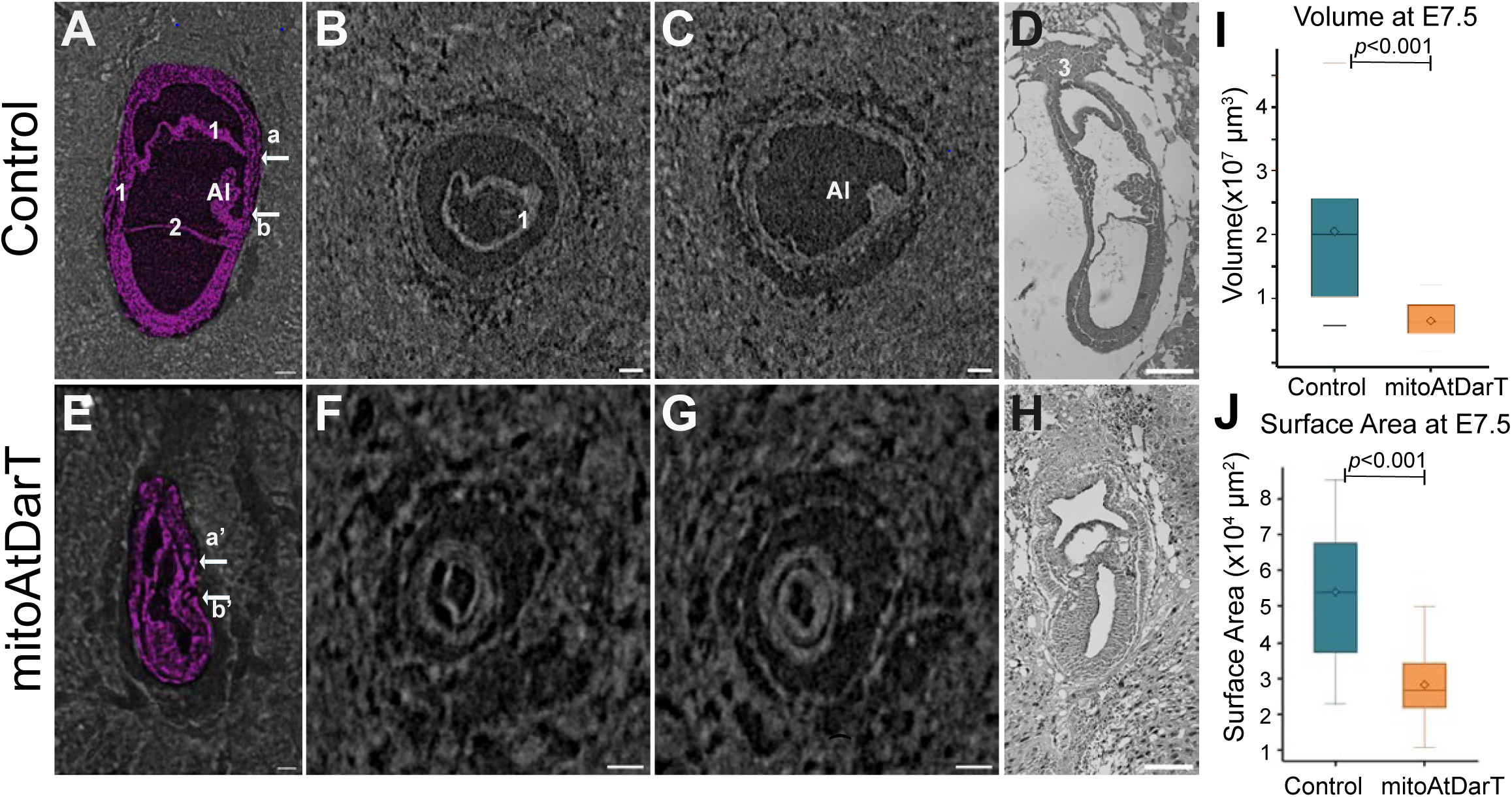
In utero characterization of R26-mitoAtDarT embryonic morphology. A. Sagittal 2D virtual section of a control (R26R-mitoAtDarT) embryo at E7.5 showing normal embryonic morphology with proper compartmentalization and lineage formation. High-resolution microCT scanning was performed at 0.7 μm voxel size. B – C. Transverse virtual sections of the control embryo corresponding to planes a and b indicated in (A). Sections highlight proper formation of extraembryonic and embryonic lineages, including the chorion (1), amnion (2), ectoplacental cone (3), and allantois (Al). D. Sagittal histological section (H&E staining) of a wild-type embryo at E7.5 showing normal embryonic architecture. E. Sagittal 2D virtual section of an R26-mitoAtDarT embryo demonstrating reduced size and disrupted morphology, including absence of major embryonic structures such as the allantois. F – G. Transverse sections corresponding to planes a′ and b′ in (E), highlighting morphological defects and size reduction compared to controls. H. Sagittal H&E-stained histological section of an R26-mitoAtDarT embryo showing severely disrupted architecture. I. Quantification of embryo volume from 3D-rendered microCT scans showing a significant reduction in volume in mitoAtDarT-expressing embryos compared to controls (*p* < 0.0001, Mann-Whitney U test) J. Surface area measurements from the same 3D renderings also reveal a significant decrease in surface area in mutant embryos (*p* < 0.0001). Control embryos were derived from R26R-mitoAtDarT × B6ND2 crosses. Whole-uterus processing and scanning precluded genotyping or GFP-based identification Scale bars: A–C, E–G, 50 μm; D, H, 100 μm. Labels: 1 – Chorion; 2 – Amnion; 3 – Ectoplacental cone; Al – Allantois. n=24 R26-mitoAtDarT embryos and 27 R26R-mitoAtDarT (control).

A consistent feature of the E7.5 phenotype was the failure to form a morphologically distinct allantois. This structure normally emerges from the posterior region in close association with the primitive streak during gastrulation (Downs et al., 2004; Downs et al., 2009). The allantois was absent from mitoAtDarT embryos despite the continued presence of other recognisable embryonic and extraembryonic structures. We therefore examined this phenotype in greater detail (see below) to determine whether it reflected a general failure of gastrulation or a more specific defect in posterior mesodermal development.

Together, these observations suggest that mitoAtDarT embryos initiate post-implantation development but progressively fail to maintain normal growth and morphogenesis. The phenotype is already detectable at E6.5, becomes markedly more severe by E7.5, and ultimately culminates in developmental arrest and resorption.

### mitoAtDarT expression is associated with reduced proliferation rather than widespread apoptosis

The reduced size of R26-mitoAtDarT embryos at E6.5 and E7.5 suggested that mitochondrial dysfunction might impair embryonic growth through reduced cell proliferation, increased cell death, or both. We therefore first examined proliferation at E7.5 using EdU incorporation to identify cells in S phase (Salic and Mitchison, 2008). Control embryos showed widespread EdU incorporation across embryonic tissues, consistent with the rapid proliferation characteristic of gastrulation (MacAuley et al., 1993). In contrast, R26-mitoAtDarT embryos contained significantly fewer EdU-positive cells (Fig. 4A, B), indicating reduced proliferative activity. To assess this independently, we examined phospho-histone H3 (pHH3), a marker of cells in mitosis (Hendzel et al., 1997). Consistent with the EdU data, the proportion of pHH3-positive cells was also reduced in R26-mitoAtDarT embryos (Supplementary Fig. 6). Together, these findings indicate that mitoAtDarT expression is associated with reduced progression through mitosis.

**Figure 4.**
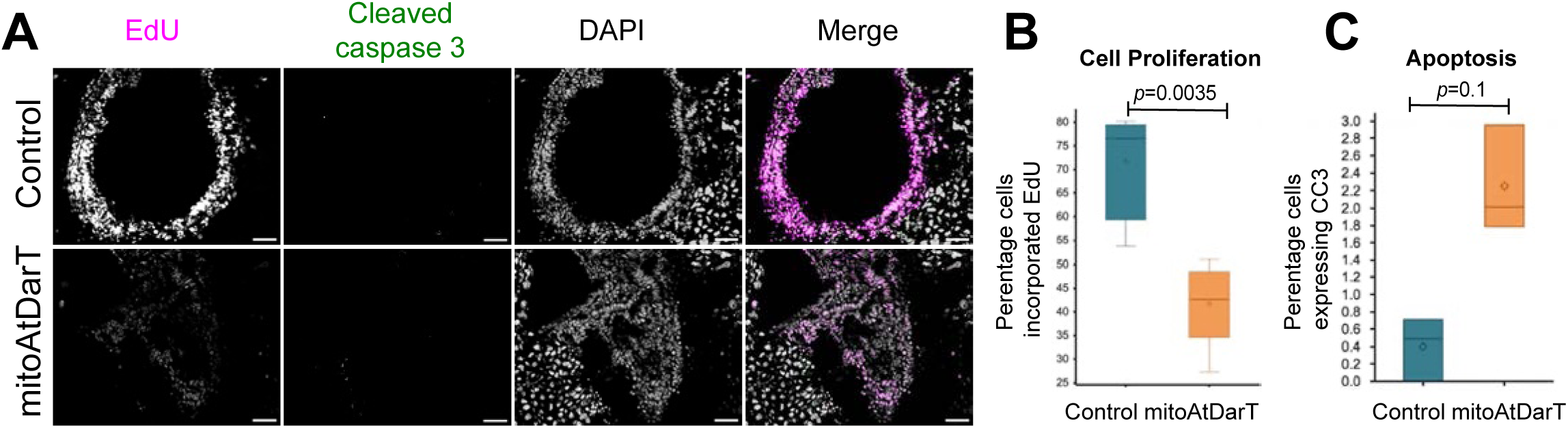
MitoAtDarT expression reduces proliferation without increasing apoptosis. A. Sagittal sections of E7.5 embryos stained for EdU (S-phase marker, magenta) and cleaved caspase-3 (apoptosis marker, green). Control (R26R-mitoAtDarT) embryos show widespread EdU incorporation, indicating active proliferation. In contrast, R26-mitoAtDarT embryos exhibit markedly reduced EdU-positive cells and signal intensity, consistent with decreased proliferation. Few cleaved caspase-3-positive cells (arrowheads). DAPI staining showing no apparent nuclear fragmentation. B. Quantification of EdU-positive cells relative to total DAPI-stained cells shows a >30% reduction in proliferation in R26-mitoAtDarT embryos compared to controls. C. Quantification of cleaved caspase-3-positive cells reveals a low (∼2%) and statistically insignificant increase in apoptosis in mutants. Scale bars, 20 μm. n = 5 R26-mitoAtDarT embryos and 4 R26R-mitoAtDarT (control).

We next asked whether increased apoptosis also contributed to the growth defect. Immunostaining for cleaved caspase-3 revealed only a small number of apoptotic cells in R26-mitoAtDarT embryos, with no evidence of widespread cell death or marked nuclear fragmentation (Fig. 4A, C). Thus, although apoptosis may contribute modestly to the phenotype, it does not appear sufficient to account for the pronounced reduction in embryo size.

These observations suggest that the growth retardation of R26-mitoAtDarT embryos is associated predominantly with reduced proliferation rather than extensive apoptosis.

Importantly, the preservation of cell viability despite impaired growth provided a basis for examining whether specific developmental programmes were also disrupted.

### mitoAtDarT expression impairs normal allantoic morphogenesis from posterior mesoderm

The absence of a morphologically distinct allantois in R26-mitoAtDarT embryos led us to ask whether this reflected a broader failure of germ-layer formation or a more specific defect in posterior mesoderm development. We therefore analysed the distribution of lineage markers in E7.5 embryos, focusing first on Brachyury (T), which is expressed in the primitive streak and nascent mesoderm during gastrulation and subsequently becomes restricted within the developing allantois to a proximal midline domain associated with the primitive streak– allantois axis (Inman and Downs, 2006b; Kispert and Herrmann, 1994; Wilkinson et al., 1990).

In control embryos, Brachyury showed the expected posterior and midline distribution, encompassing the primitive streak, axial mesoderm, and proximal allantoic region (Fig. 5A). In R26-mitoAtDarT embryos, however, the Brachyury-positive domain was expanded posteriorly and extended further into the extraembryonic mesoderm despite the absence of a morphologically distinct allantois. During normal allantoic development, Brachyury becomes spatially restricted as the allantois elongates, with more distal allantoic cells lacking Brachyury expression (Inman and Downs, 2006a; Inman and Downs, 2006b).

**Figure 5.**
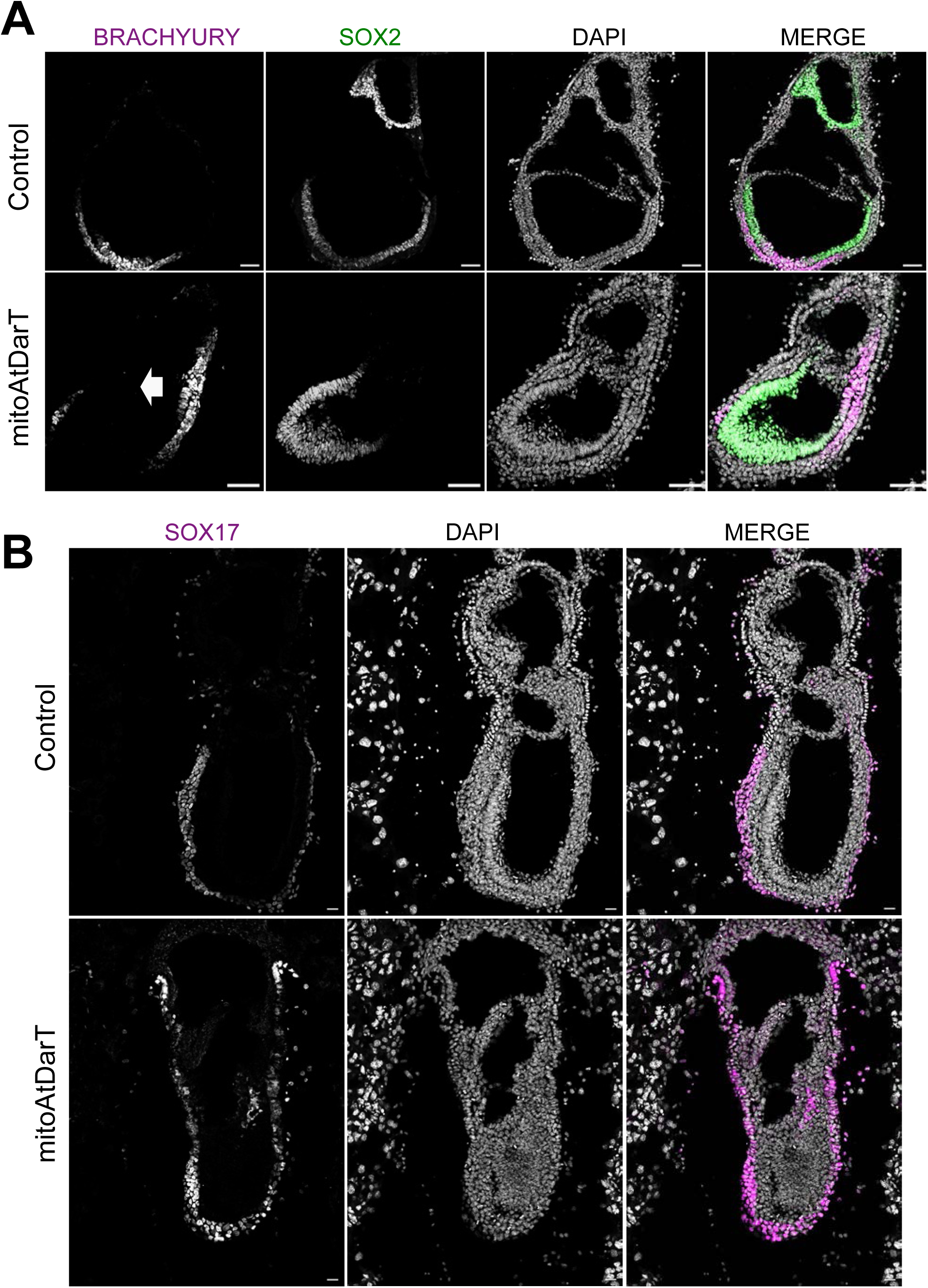
MitoAtDarT disrupts mesoderm patterning at E7.5. A. Immunofluorescence staining of Brachyury (magenta), SOX2 (green), and nuclei (DAPI, gray) in sagittal sections of E7.5 embryos. In control (R26R-mitoAtDarT) embryos, SOX2 SOX2 marks the epiblast and neural ectoderm, while Brachyury expression is restricted to mesodermal cells within the notochord region. In R26-mitoAtDarT embryos, Brachyury expression is aberrantly extended into extraembryonic regions. SOX2 expression remains intact. Scale bars, 20 μm. n = 5 embryos per genotype. B. Coronal sections of E7.5 embryos stained for SOX17 (magenta) and nuclei (DAPI, gray). In control embryos, SOX17 expression is localized to the visceral endoderm with sparse positive cells in the allantois. In R26-mitoAtDarT embryos, SOX17-positive cells surround the embryo as expected but appears abnormally enriched in the allantoic bud region. Scale bars, 20 μm. n = 3 embryos per genotype.

The extended Brachyury-positive domain in R26-mitoAtDarT embryos therefore suggests that posterior mesoderm is generated but fails to undergo the normal spatial resolution of Brachyury expression associated with allantoic development.

We next examined whether other major lineage-associated compartments were also perturbed. SOX2-positive cells, marking epiblast-derived ectodermal populations at this stage (Wood and Episkopou, 1999), remained readily detectable in R26-mitoAtDarT embryos and showed a broadly comparable distribution to controls (Fig. 5A). Similarly, SOX17-positive cells remained distributed around the outer layer of mutant embryos, indicating preservation of SOX17-positive endodermal populations (Kanai-Azuma et al., 2002), although these cells appeared enriched around the posterior/allantoic-bud region in some embryos (Fig. 5B).

Thus, despite the pronounced morphological abnormalities of R26-mitoAtDarT embryos, major SOX2-, SOX17- and Brachyury-positive cellular compartments remained recognisable.

These observations suggest that the allantoic defect cannot be explained simply by a general failure to establish the major embryonic lineages. Instead, the coexistence of an expanded Brachyury-positive posterior domain with failure of allantoic outgrowth may point to an abnormality in the subsequent organisation or progression of posterior mesoderm. One possibility is that cells enter a Brachyury-positive mesodermal state but fail to progress normally through the developmental programme required for allantois formation. This would be consistent with the persistence of incompletely resolved lineage identities observed in early R26-mitoAtDarT blastocysts and raises the possibility that mitochondrial dysfunction compromises the timely progression of cells between developmental states.

### mitoAtDarT expression is accompanied by metabolic remodelling during gastrulation

Given the pronounced mitochondrial dysfunction in R26-mitoAtDarT embryos, we next asked whether this was accompanied by metabolic changes that could influence developmental signalling during gastrulation. Metabolic state is increasingly recognised as an important regulator of signalling and morphogenesis during embryonic development (Bulusu et al., 2017; Cao et al., 2024; Miyazawa et al., 2022). In particular, glycolytic activity can modulate Wnt and Nodal signalling, pathways that are themselves essential for primitive streak formation and mesoderm development (Conlon et al., 1994; Liu et al., 1999; Stapornwongkul et al., 2025). We therefore examined the metabolic state of E7.5 R26-mitoAtDarT embryos.

We first analysed pyruvate kinase M2 (PKM2), a key glycolytic enzyme that catalyses the final step of glycolysis. PKM2 immunoreactivity was elevated in R26-mitoAtDarT embryos compared with controls, with particularly strong enrichment in the posterior region containing the primitive streak and nascent mesoderm (Fig. 6A, B). This regional increase was notable given that glycolytic activity is spatially regulated within posterior embryonic tissues and can influence developmental signalling and mesodermal patterning (Bulusu et al., 2017; Miyazawa et al., 2022), and given the abnormal persistence and distribution of Brachyury-positive posterior mesoderm observed at these stages.

**Figure 6.**
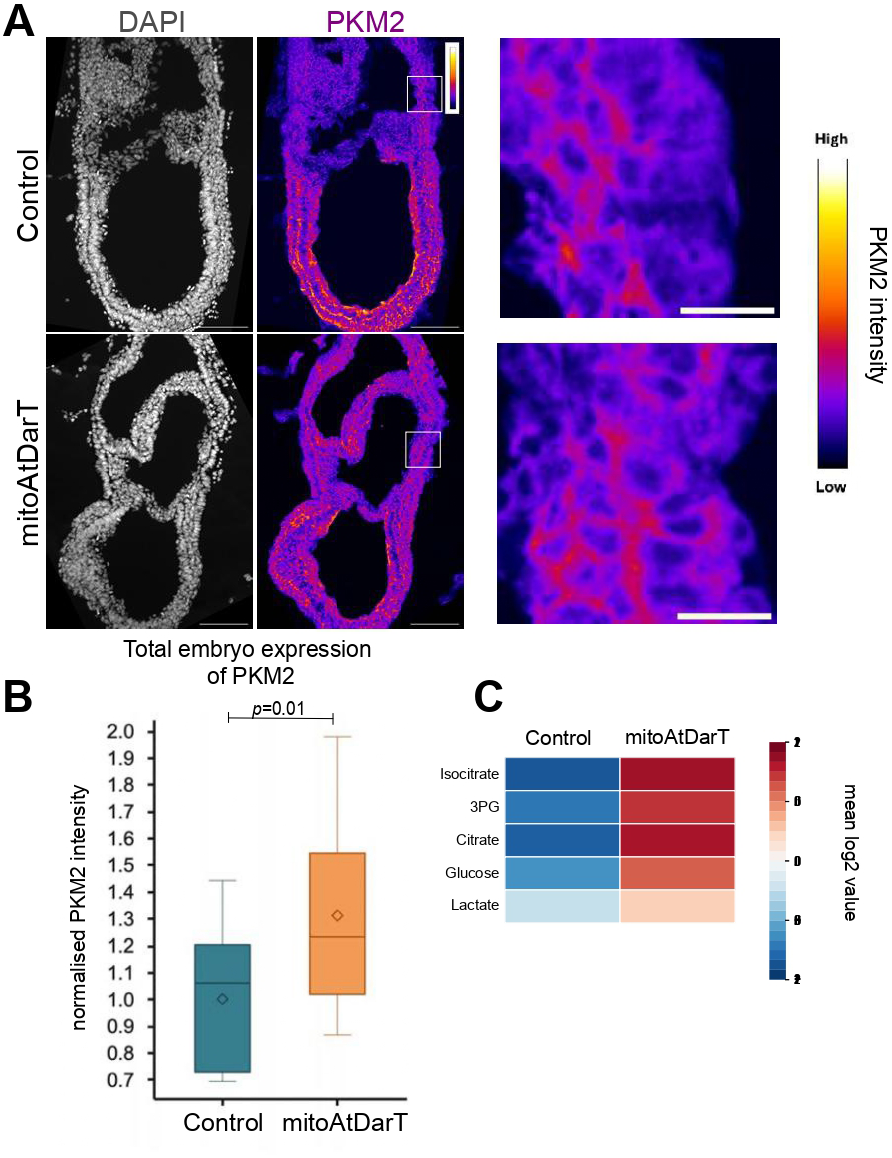
Metabolic remodelling and glycolysis upregulation in R26-mitoAtDarT embryos. A. Representative Immunofluorescence images of E7.5 control (R26R-mitoAtDarT) embryos and R26-mitoAtDarT stained for nuclei (DAPI, gray) and PKM2 a marker for glycolysis (fire LUT). PKM2 intensity is increased in R26-mitoAtDarT compared to control. The right images are higher magnification of the regions marked with the grey squares from the PKM2 channel. B. Quantification of PKM2 fluorescent intensity in control and R26-mitoAtDarT embryos. Data are presented from n = 5 embryos per genotype. P=0.001. C. Heat map of relative metabolite abundance in the control R26R-mitoAtDarT and R26-mitoAtDarT embryos measured using LC-MS. Data were collected from three biological replicates (n=3). Only statistically significant (P<0.05) metabolites are shown. Isocitrate, 3-phosphoglycerate (3PG), citrate, glucose, and Lactate showed upregulation in mitoAtDarT-expressing embryos.

To assess metabolic changes more broadly, we performed targeted mass-spectrometric profiling of glycolytic, pentose phosphate pathway and TCA-cycle metabolites. Although several individual metabolites did not reach statistical significance, the overall profile showed a tendency towards increased abundance of multiple glucose-associated metabolites. These included the glycolytic products 3-phosphoglycerate and lactate as well as an accumulation of isocitrate and citrate (Fig. 6C).

Thus, mitochondrial dysfunction in R26-mitoAtDarT embryos is accompanied by remodelling of glucose-associated metabolism during gastrulation. The coincidence of these metabolic changes with abnormal posterior mesoderm development is notable given evidence that glycolytic state can modulate Wnt, Nodal and FGF signalling and thereby influence mesodermal patterning and developmental progression (Miyazawa et al., 2022; Oginuma et al., 2017; Stapornwongkul et al., 2025). Although our data do not establish a direct causal relationship between altered metabolism and the posterior mesoderm phenotype, they suggest that mitochondrial dysfunction changes the metabolic environment in which these developmental programmes operate. These changes could contribute to the failure of Brachyury-positive cells to undergo normal progression towards an allantoic mesodermal state, providing a potential link between mitochondrial function, metabolic state and developmental progression.

## DISCUSSION

In this study, we establish mitoAtDarT as a conditional system for perturbing mitochondrial function during mammalian development and show that its early activation disrupts embryonic progression. R26-mitoAtDarT embryos form blastocysts and implant, but subsequently exhibit reduced growth, abnormal morphogenesis and embryonic lethality. At two distinct developmental stages, the phenotype is associated with persistence of incompletely resolved cellular states: CDX2/SOX2 double-positive cells are increased in blastocysts, while Brachyury-positive posterior mesoderm persists despite failure of allantoic morphogenesis. These observations raise the possibility that mitochondrial function contributes not only to embryonic growth but also to the timely progression of cells between developmental states.

MitoAtDarT provides an approach complementary to genetic models that disrupt mitochondrial genome maintenance, such as loss of Tfam or Polg, which result in severe mtDNA depletion and early embryonic lethality (Hance et al., 2005; Larsson et al., 1998). Mitochondrially targeted DarT has previously been shown to damage mtDNA and impair mitochondrial function in yeast (Dua et al., 2022). In R26-mitoAtDarT embryos, AtDarT localised efficiently to mitochondria and was associated with reduced membrane potential, reduced mitochondrial area and altered mitochondrial morphology. We detected no increase in nuclear γH2AX, arguing against a substantial nuclear double-strand-break response under these conditions. Direct measurement of mtDNA lesions will nevertheless be required to establish the extent and distribution of mitochondrial genome damage in the mouse embryo. An important feature of the R26R-mitoAtDarT model is its conditionality, which should ultimately allow mitochondrial requirements to be examined with greater temporal and lineage specificity.

The earliest developmental phenotype was apparent during blastocyst formation. Although R26-mitoAtDarT blastocysts contained fewer cells, the relative representation of clearly CDX2-positive and SOX2-positive populations was broadly retained. The more striking difference was an increased population co-expressing both factors. Rather than indicating a simple shift in allocation between trophectoderm and inner cell mass, this is consistent with delayed or incomplete resolution of lineage-associated transcriptional states. Metabolism provides a plausible route through which mitochondrial dysfunction could affect this process. Glucose metabolism has been shown to regulate trophectoderm specification through YAP–TEAD4-dependent mechanisms, linking metabolic state directly to the transcriptional machinery controlling the first lineage decision (Chi et al., 2020; Nishioka et al., 2009). We have not tested whether this pathway is altered in R26-mitoAtDarT embryos, but these observations provide a precedent for metabolic state influencing the resolution of TE and ICM identities.

A related but more spatially restricted phenotype becomes apparent during gastrulation. Despite severe growth retardation, R26-mitoAtDarT embryos showed SOX2- and SOX17-positive populations and generated Brachyury-positive mesoderm. The phenotype therefore does not reflect a general failure to establish the major germ-layers.

Instead, the Brachyury-positive domain was expanded within the posterior region, while a morphologically distinct allantois failed to form. This is important because Brachyury expression alone does not indicate successful progression through an allantoic programme. During normal development, Brachyury becomes spatially restricted within the proximal midline of the developing allantois rather than being uniformly maintained throughout the elongating structure, and Brachyury function is required for normal allantoic elongation and organisation (Inman and Downs, 2006a; Inman and Downs, 2006b). BMP4 signalling is required for formation and subsequent development of the allantois (Fujiwara et al., 2001; Lawson et al., 1999), and may suggest a possible pathway that mitochondria could intersect with in the development of the allantois. Thus, the presence of an extended Brachyury-positive posterior domain in R26-mitoAtDarT embryos indicates that posterior mesoderm is generated but does not undergo the normal spatial and morphogenetic progression associated with allantois formation.

This interpretation is relevant given increasing evidence that metabolism participates directly in coordinating fate specification and morphogenesis. In the gastrulating mouse embryo, spatially regulated glucose metabolism promotes epiblast fate acquisition and guides mesoderm migration through developmental signalling (Cao et al., 2024). Stem-cell-derived embryo models further show that glycolytic activity regulates Wnt and Nodal signalling and germ-layer allocation, while the balance between glycolysis and oxidative phosphorylation influences subsequent lineage composition and morphology (Stapornwongkul et al., 2025; Villaronga-Luque et al., 2025). R26-mitoAtDarT embryos themselves showed increased PKM2 expression, particularly posteriorly. This does suggest that mitochondrial dysfunction is accompanied by remodelling of glucose-associated metabolism. Altered metabolic state could therefore modify the signalling, redox or chromatin environment in which posterior mesoderm develops (Cao et al., 2024; Stapornwongkul et al., 2025; Zhao et al., 2021).

Testing this possibility will require direct manipulation of these metabolic pathways. A limitation of the present study is that mitoAtDarT is activated from the earliest stages of embryogenesis. Development is already altered at the blastocyst stage, and growth retardation is evident by E6.5, making it difficult to determine whether later abnormalities reflect direct mitochondrial requirements within particular lineages or consequences of earlier defects. The current data therefore cannot distinguish between a broadly permissive requirement for mitochondrial function and selective sensitivity of particular developmental transitions. This distinction can be addressed using the conditional R26R-mitoAtDarT allele to restrict mitochondrial perturbation temporally or to defined progenitor populations. Such experiments should separate embryo-wide effects from lineage-specific requirements and reveal whether different developmental programmes depend on distinct aspects of mitochondrial activity.

Overall, our findings show that early mammalian embryos can establish several major lineage-associated programmes despite mitochondrial dysfunction, but subsequent growth, lineage resolution and morphogenesis are compromised. Rather than acting solely as a source of bioenergetic support, mitochondria may therefore form part of the regulatory environment that allows developmental programmes to unfold in the appropriate sequence. Defining how mitochondrial state interacts with signalling and transcriptional networks should help establish whether this represents a common feature of developmental transitions or a particular vulnerability of selected embryonic lineages.

## ACKNOWLEDGEMENTS

This work was supported by the Department of Atomic Energy, Government of India, Project Identification No. RTI 4006 and HFSP RPG to AB (RGP0038/2021-102). We thank the NCBS Mouse Genome Engineering Facility, especially Dr Aurelie Jory for generating the mouse and Dr Mahesh Sahare for blastocyst collection. We thank Mr. Sunil Prabhakar from the MicroCT facility for help in establishing the imaging protocol and developing the analysis pipeline. We would also like to thank Dr. Venkatesan Iyer from the CIFF facility for his assistance with imaging. We are grateful to Dr. Nitish Dua for discussion and Dr. Akshaya Seshadri for validating the activity of AtDarT in yeast cells. We thank the members of the Ear Lab at NCBS for their feedback and discussion.

## AUTHOR CONTRIBUTIONS

Conceptualization: R.K.L., A.B., MA; Methodology: M.A., N.G., G.P., M.H.; Funding acquisition: R.K.L., A.B.; Project administration: R.K.L.; Supervision: R.K.L., A.B.; Writing – original draft: M.A.; Writing – review & editing: R.K.L.

## AI STATEMENT

During the preparation of this work, the author(s) used ChatGPT for Proofreading the manuscript. The author(s) reviewed and edited the output as needed and take full responsibility for the content of the published article.

## REFERENCES

1. Beddington, R.S., and E.J. Robertson. 1999. Axis development and early asymmetry in mammals. Cell. 96:195–209.

2. Bulusu, V., N. Prior, M.T. Snaebjornsson, A. Kuehne, K.F. Sonnen, J. Kress, F. Stein, C. Schultz, U. Sauer, and A. Aulehla. 2017. Spatiotemporal Analysis of a Glycolytic Activity Gradient Linked to Mouse Embryo Mesoderm Development. Dev Cell. 40:331–341 e334.

3. Cao, D., J. Bergmann, L. Zhong, A. Hemalatha, C. Dingare, T. Jensen, A.L. Cox, V. Greco, B. Steventon, and B. Sozen. 2024. Selective utilization of glucose metabolism guides mammalian gastrulation. Nature. 634:919–928.

4. Chakrabarty, R.P., and N.S. Chandel. 2021. Mitochondria as Signaling Organelles Control Mammalian Stem Cell Fate. Cell stem cell. 28:394–408.

5. Chandel, N.S. 2014. Mitochondria as signaling organelles. BMC biology. 12:34.

6. Chi, F., M.S. Sharpley, R. Nagaraj, S.S. Roy, and U. Banerjee. 2020. Glycolysis-Independent Glucose Metabolism Distinguishes TE from ICM Fate during Mammalian Embryogenesis. Dev Cell. 53:9–26 e24.

7. Christodoulou, N., A. Weberling, D. Strathdee, K.I. Anderson, P. Timpson, and M. Zernicka-Goetz. 2019. Morphogenesis of extra-embryonic tissues directs the remodelling of the mouse embryo at implantation. Nature communications. 10:3557.

8. Chu, V.T., T. Weber, R. Graf, T. Sommermann, K. Petsch, U. Sack, P. Volchkov, K. Rajewsky, and R. Kuhn. 2016. Efficient generation of Rosa26 knock-in mice using CRISPR/Cas9 in C57BL/6 zygotes. BMC Biotechnol. 16:4.

9. Conlon, F.L., K.M. Lyons, N. Takaesu, K.S. Barth, A. Kispert, B. Herrmann, and E.J. Robertson. 1994. A primary requirement for nodal in the formation and maintenance of the primitive streak in the mouse. Development. 120:1919–1928.

10. Downs, K.M., E.R. Hellman, J. McHugh, K. Barrickman, and K.E. Inman. 2004. Investigation into a role for the primitive streak in development of the murine allantois. Development. 131:37–55.

11. Downs, K.M., K.E. Inman, D.X. Jin, and A.C. Enders. 2009. The Allantoic Core Domain: new insights into development of the murine allantois and its relation to the primitive streak. Dev Dyn. 238:532–553.

12. Dua, N., A. Seshadri, and A. Badrinarayanan. 2022. DarT-mediated mtDNA damage induces dynamic reorganization and selective segregation of mitochondria. J Cell Biol. 221.

13. Ermakova, O., T. Orsini, A. Gambadoro, F. Chiani, and G.P. Tocchini-Valentini. 2018. Three-dimensional microCT imaging of murine embryonic development from immediate post-implantation to organogenesis: application for phenotyping analysis of early embryonic lethality in mutant animals. Mammalian genome : official journal of the International Mammalian Genome Society. 29:245–259.

14. Facucho-Oliveira, J.M., and J.C. St John. 2009. The relationship between pluripotency and mitochondrial DNA proliferation during early embryo development and embryonic stem cell differentiation. Stem Cell Rev Rep. 5:140–158.

15. Fujiwara, T., N.R. Dunn, and B.L. Hogan. 2001. Bone morphogenetic protein 4 in the extraembryonic mesoderm is required for allantois development and the localization and survival of primordial germ cells in the mouse. Proc Natl Acad Sci U S A. 98:13739–13744.

16. Gokbuget, D., and R. Blelloch. 2019. Epigenetic control of transcriptional regulation in pluripotency and early differentiation. Development. 146.

17. Hance, N., M.I. Ekstrand, and A. Trifunovic. 2005. Mitochondrial DNA polymerase gamma is essential for mammalian embryogenesis. Hum Mol Genet. 14:1775–1783.

18. Harvey, A., G. Caretti, V. Moresi, A. Renzini, and S. Adamo. 2019. Interplay between Metabolites and the Epigenome in Regulating Embryonic and Adult Stem Cell Potency and Maintenance. Stem cell reports. 13:573–589.

19. Houghton, F.D., J.G. Thompson, C.J. Kennedy, and H.J. Leese. 1996. Oxygen consumption and energy metabolism of the early mouse embryo. Mol Reprod Dev. 44:476–485.

20. Inman, K.E., and K.M. Downs. 2006a. Brachyury is required for elongation and vasculogenesis in the murine allantois. Development. 133:2947–2959.

21. Inman, K.E., and K.M. Downs. 2006b. Localization of Brachyury (T) in embryonic and extraembryonic tissues during mouse gastrulation. Gene Expr Patterns. 6:783–793.

22. Jankevicius, G., A. Ariza, M. Ahel, and I. Ahel. 2016. The Toxin-Antitoxin System DarTG Catalyzes Reversible ADP-Ribosylation of DNA. Mol Cell. 64:1109–1116.

23. Kanai-Azuma, M., Y. Kanai, J.M. Gad, Y. Tajima, C. Taya, M. Kurohmaru, Y. Sanai, H. Yonekawa, K. Yazaki, P.P. Tam, and Y. Hayashi. 2002. Depletion of definitive gut endoderm in Sox17-null mutant mice. Development. 129:2367–2379.

24. Kasahara, A., S. Cipolat, Y. Chen, G.W. Dorn, 2nd, and L. Scorrano. 2013. Mitochondrial fusion directs cardiomyocyte differentiation via calcineurin and Notch signaling. Science. 342:734–737.

25. Kispert, A., and B.G. Herrmann. 1994. Immunohistochemical analysis of the Brachyury protein in wild-type and mutant mouse embryos. Dev Biol. 161:179–193.

26. Lallemand, Y., V. Luria, R. Haffner-Krausz, and P. Lonai. 1998. Maternally expressed PGK-Cre transgene as a tool for early and uniform activation of the Cre site-specific recombinase. Transgenic research. 7:105–112.

27. Larsson, N.G., J. Wang, H. Wilhelmsson, A. Oldfors, P. Rustin, M. Lewandoski, G.S. Barsh, and D.A. Clayton. 1998. Mitochondrial transcription factor A is necessary for mtDNA maintenance and embryogenesis in mice. Nat Genet. 18:231–236.

28. Lawaree, E., G. Jankevicius, C. Cooper, I. Ahel, S. Uphoff, and C.M. Tang. 2020. DNA ADP-Ribosylation Stalls Replication and Is Reversed by RecF-Mediated Homologous Recombination and Nucleotide Excision Repair. Cell reports. 30:1373–1384 e1374.

29. Lawson, K.A., N.R. Dunn, B.A. Roelen, L.M. Zeinstra, A.M. Davis, C.V. Wright, J.P. Korving, and B.L. Hogan. 1999. Bmp4 is required for the generation of primordial germ cells in the mouse embryo. Genes Dev. 13:424–436.

30. Leese, H.J., and A.M. Barton. 1984. Pyruvate and glucose uptake by mouse ova and preimplantation embryos. J Reprod Fertil. 72:9–13.

31. Liu, P., M. Wakamiya, M.J. Shea, U. Albrecht, R.R. Behringer, and A. Bradley. 1999. Requirement for Wnt3 in vertebrate axis formation. Nat Genet. 22:361–365.

32. Miroshnikova, Y.A., M.N. Shahbazi, J. Negrete, K.J. Chalut, and A. Smith. 2023. Cell state transitions: catch them if you can. Development. 150.

33. Miyazawa, H., and A. Aulehla. 2018. Revisiting the role of metabolism during development. Development. 145.

34. Miyazawa, H., M.T. Snaebjornsson, N. Prior, E. Kafkia, H.M. Hammaren, N. Tsuchida-Straeten, K.R. Patil, M. Beck, and A. Aulehla. 2022. Glycolytic flux-signaling controls mouse embryo mesoderm development. eLife. 11.

35. Morgani, S.M., and A.K. Hadjantonakis. 2020. Signaling regulation during gastrulation: Insights from mouse embryos and in vitro systems. Curr Top Dev Biol. 137:391–431.

36. Moris, N., C. Pina, and A.M. Arias. 2016. Transition states and cell fate decisions in epigenetic landscapes. Nat Rev Genet. 17:693–703.

37. Nishioka, N., K. Inoue, K. Adachi, H. Kiyonari, M. Ota, A. Ralston, N. Yabuta, S. Hirahara, R.O. Stephenson, N. Ogonuki, R. Makita, H. Kurihara, E.M. Morin-Kensicki, H. Nojima, J. Rossant, K. Nakao, H. Niwa, and H. Sasaki. 2009. The Hippo signaling pathway components Lats and Yap pattern Tead4 activity to distinguish mouse trophectoderm from inner cell mass. Dev Cell. 16:398–410.

38. Oginuma, M., P. Moncuquet, F. Xiong, E. Karoly, J. Chal, K. Guevorkian, and O. Pourquie. 2017. A Gradient of Glycolytic Activity Coordinates FGF and Wnt Signaling during Elongation of the Body Axis in Amniote Embryos. Dev Cell. 40:342–353 e310.

39. Posfai, E., S. Petropoulos, F.R.O. de Barros, J.P. Schell, I. Jurisica, R. Sandberg, F. Lanner, and J. Rossant. 2017. Position- and Hippo signaling-dependent plasticity during lineage segregation in the early mouse embryo. eLife. 6.

40. Rizzuto, R., D. De Stefani, A. Raffaello, and C. Mammucari. 2012. Mitochondria as sensors and regulators of calcium signalling. Nat Rev Mol Cell Biol. 13:566–578.

41. Rossant, J., and P.P. Tam. 2009. Blastocyst lineage formation, early embryonic asymmetries and axis patterning in the mouse. Development. 136:701–713.

42. Scaduto, R.C., Jr., and L.W. Grotyohann. 1999. Measurement of mitochondrial membrane potential using fluorescent rhodamine derivatives. Biophys J. 76:469–477.

43. Schindelin, J., I. Arganda-Carreras, E. Frise, V. Kaynig, M. Longair, T. Pietzsch, S. Preibisch, C. Rueden, S. Saalfeld, B. Schmid, J.Y. Tinevez, D.J. White, V. Hartenstein, K. Eliceiri, P. Tomancak, and A. Cardona. 2012. Fiji: an open-source platform for biological-image analysis. Nature methods. 9:676–682.

44. Sharpley, M.S., F. Chi, J.T. Hoeve, and U. Banerjee. 2021. Metabolic plasticity drives development during mammalian embryogenesis. Dev Cell. 56:2329–2347 e2326.

45. Stapornwongkul, K.S., E. Hahn, P. Polinski, L. Salamo Palau, K. Arato, L. Yao, K. Williamson, N. Gritti, K. Anlas, M. Osuna Lopez, K.R. Patil, I. Heemskerk, M. Ebisuya, and V. Trivedi. 2025. Glycolytic activity instructs germ layer proportions through regulation of Nodal and Wnt signaling. Cell stem cell. 32:744–758 e747.

46. Stern, S., J.D. Biggers, and E. Anderson. 1971. Mitochondria and early development of the mouse. J Exp Zool. 176:179–191.

47. Strumpf, D., C.A. Mao, Y. Yamanaka, A. Ralston, K. Chawengsaksophak, F. Beck, and J. Rossant. 2005. Cdx2 is required for correct cell fate specification and differentiation of trophectoderm in the mouse blastocyst. Development. 132:2093–2102.

48. Tam, P.P., and D.A. Loebel. 2007. Gene function in mouse embryogenesis: get set for gastrulation. Nat Rev Genet. 8:368–381.

49. Villaronga-Luque, A., R.G. Savill, N. Lopez-Anguita, A. Bolondi, S. Garai, S.I. Gassaloglu, R. Rouatbi, K. Schmeisser, A. Poddar, L. Bauer, T. Alves, S. Traikov, J. Rodenfels, T. Chavakis, A. Bulut-Karslioglu, and J.V. Veenvliet. 2025. Integrated molecular-phenotypic profiling reveals metabolic control of morphological variation in a stem-cell-based embryo model. Cell stem cell. 32:759–777 e713.

50. Wanet, A., T. Arnould, M. Najimi, and P. Renard. 2015. Connecting Mitochondria, Metabolism, and Stem Cell Fate. Stem cells and development. 24:1957–1971.

51. Wicklow, E., S. Blij, T. Frum, Y. Hirate, R.A. Lang, H. Sasaki, and A. Ralston. 2014. HIPPO pathway members restrict SOX2 to the inner cell mass where it promotes ICM fates in the mouse blastocyst. PLoS Genet. 10:e1004618.

52. Wilkinson, D.G., S. Bhatt, and B.G. Herrmann. 1990. Expression pattern of the mouse T gene and its role in mesoderm formation. Nature. 343:657–659.

53. Wood, H.B., and V. Episkopou. 1999. Comparative expression of the mouse Sox1, Sox2 and Sox3 genes from pre-gastrulation to early somite stages. Mech Dev. 86:197–201.

54. Zhao, J., K. Yao, H. Yu, L. Zhang, Y. Xu, L. Chen, Z. Sun, Y. Zhu, C. Zhang, Y. Qian, S. Ji, H. Pan, M. Zhang, J. Chen, C. Correia, T. Weiskittel, D.W. Lin, Y. Zhao, S. Chandrasekaran, X. Fu, D. Zhang, H.Y. Fan, W. Xie, H. Li, Z. Hu, and J. Zhang. 2021. Metabolic remodelling during early mouse embryo development. Nat Metab. 3:1372–1384.

