## Supplementary for "Disruption of mitochondrial integrity impairs early lineage specification and embryonic development in the novel conditional MitoAtDarT mouse model"

### Supplementary Figure 1

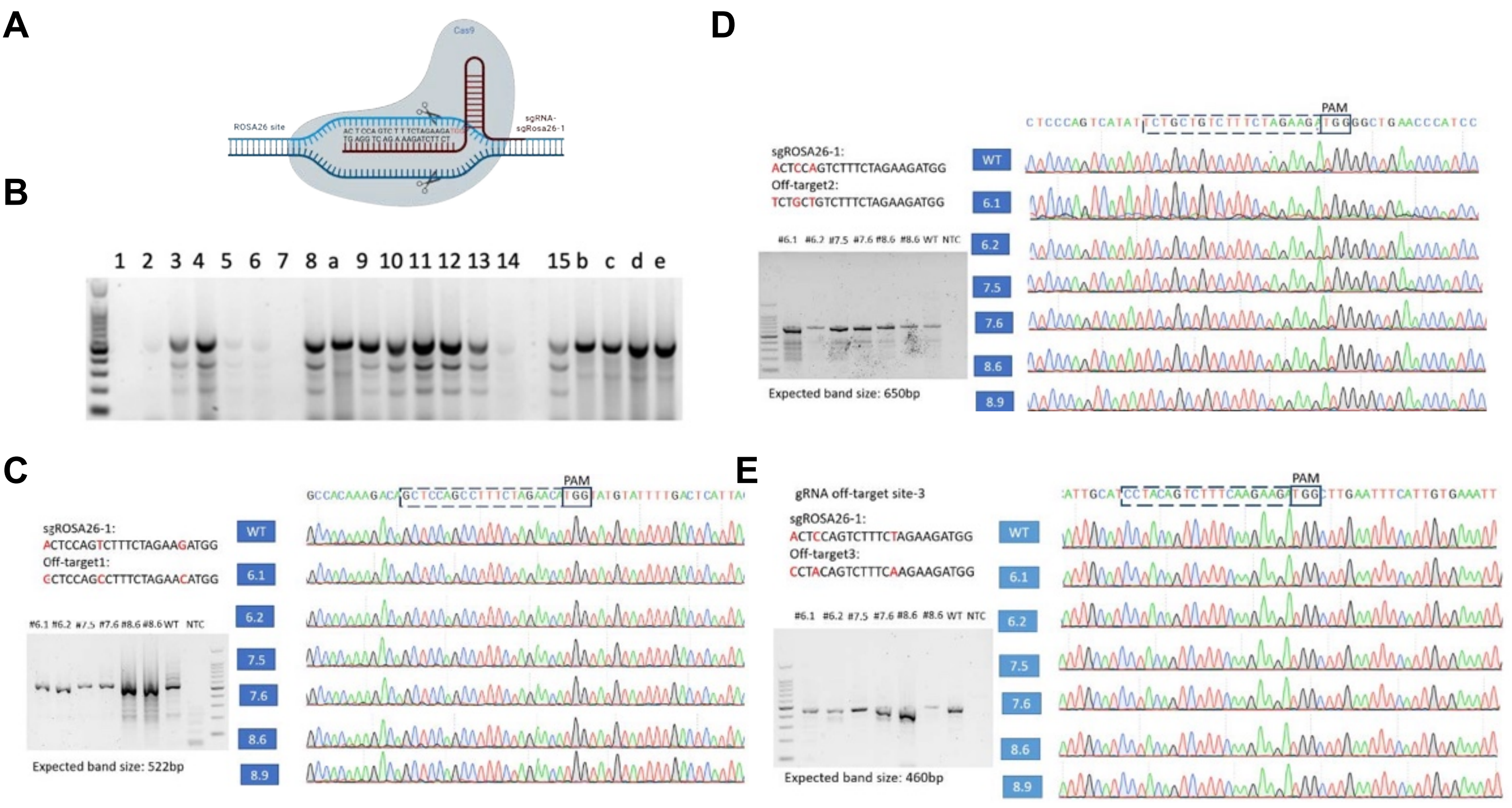

### Supplementary Figure 2

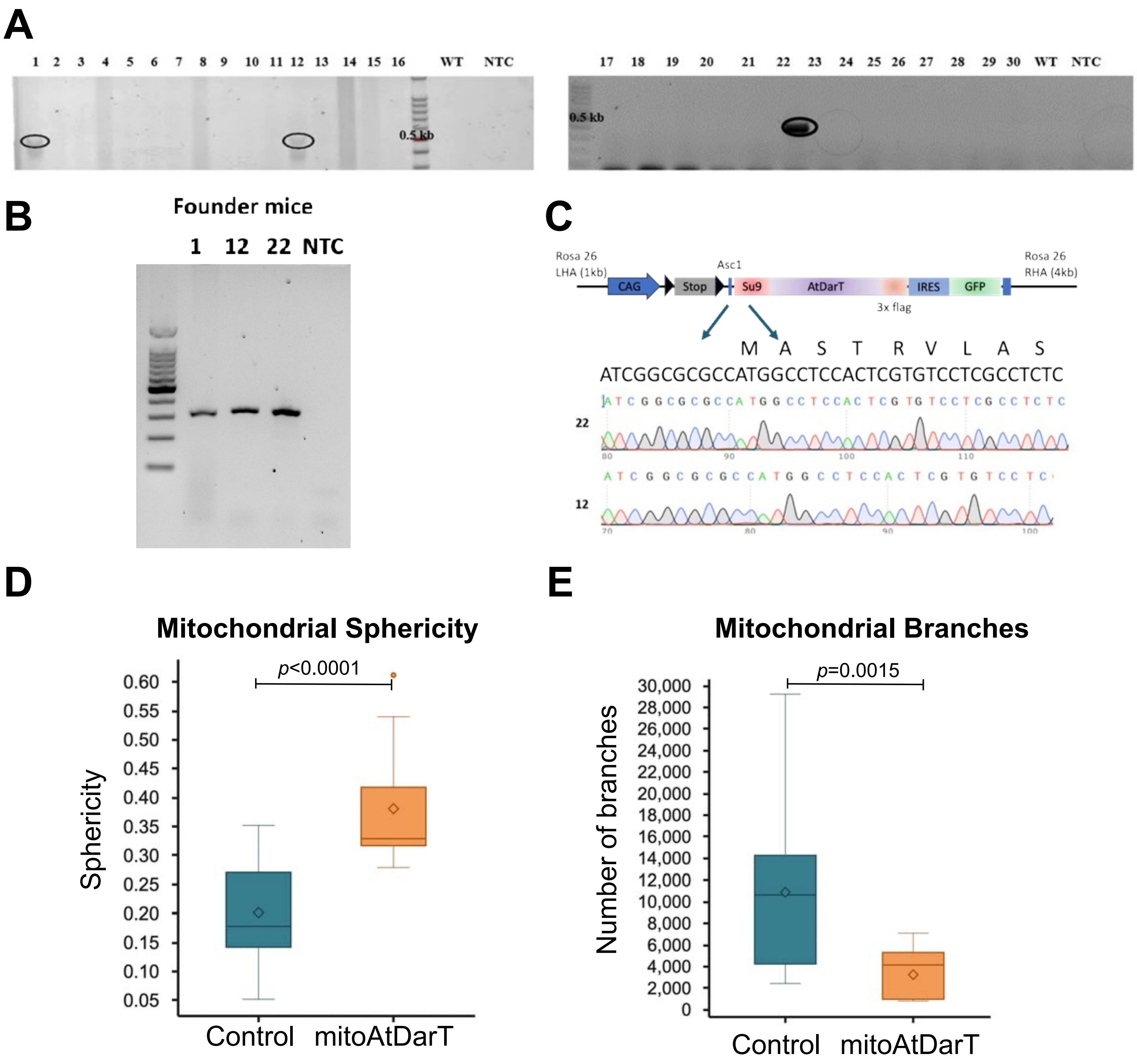

**Supplementary Figure 3**

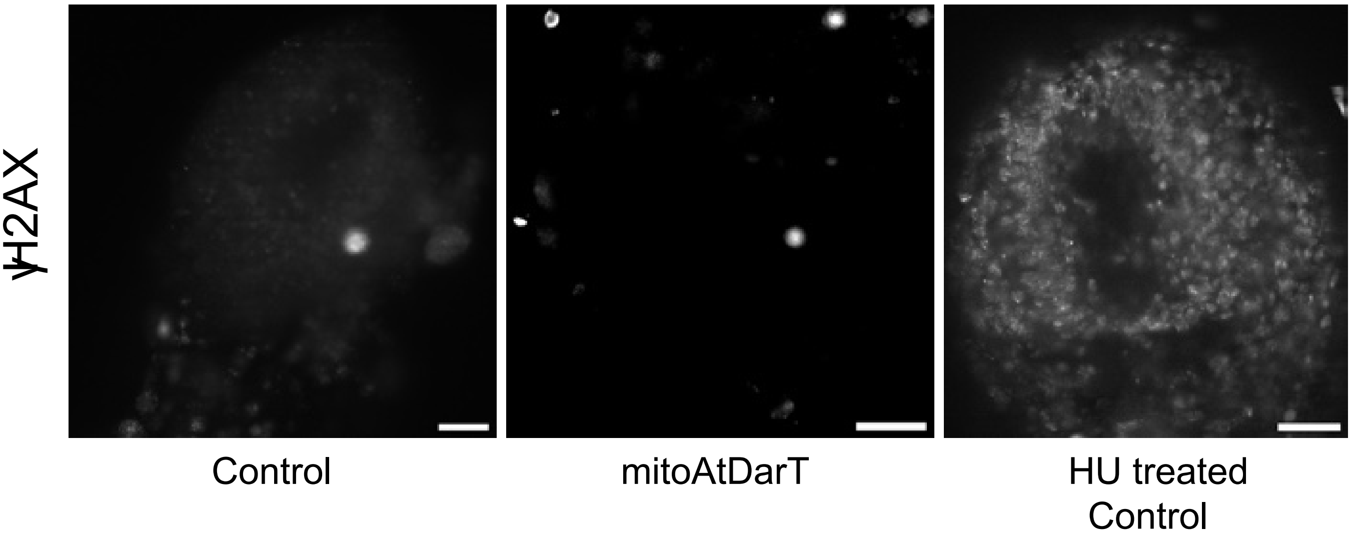

### Supplementary Figure 4

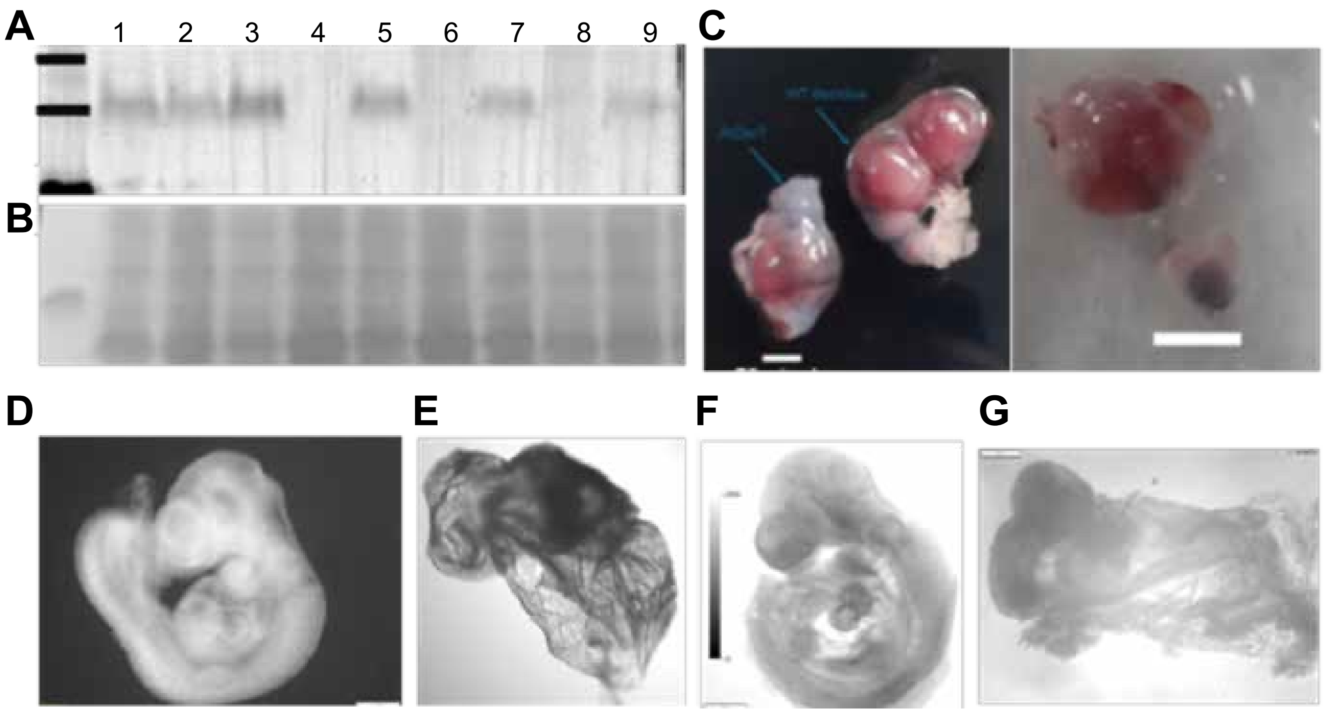

#### Supplementary Figure 5

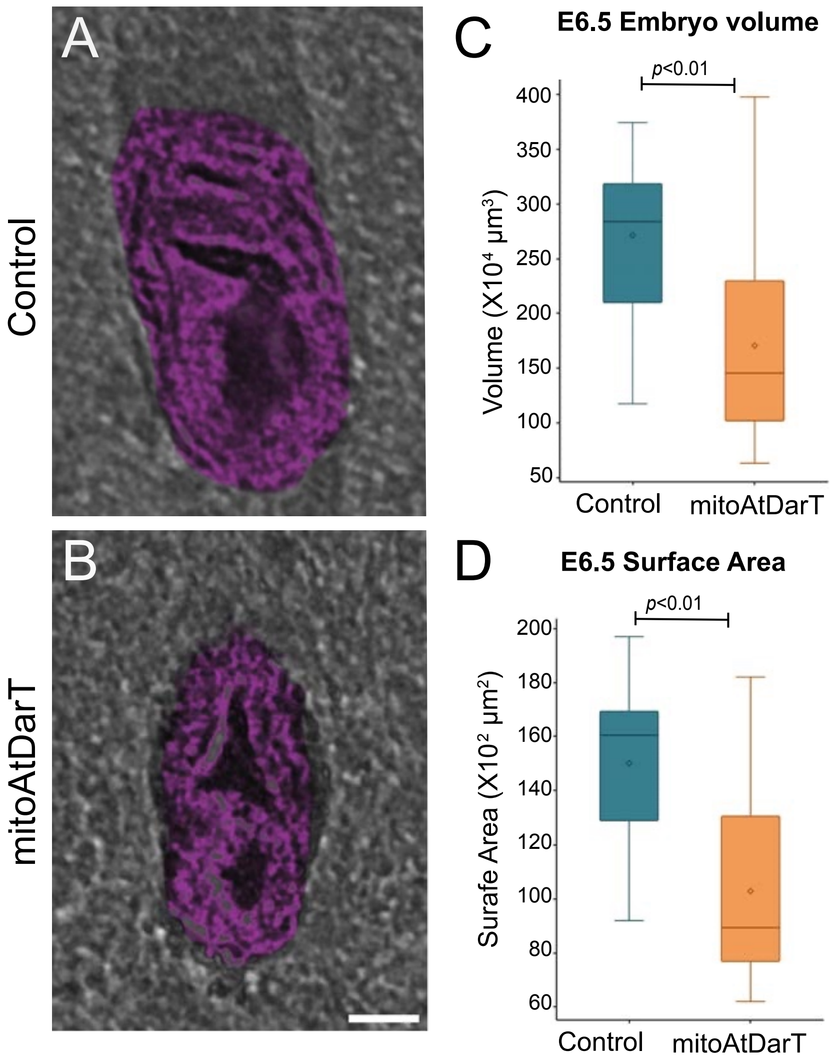

#### Supplementary Figure 6

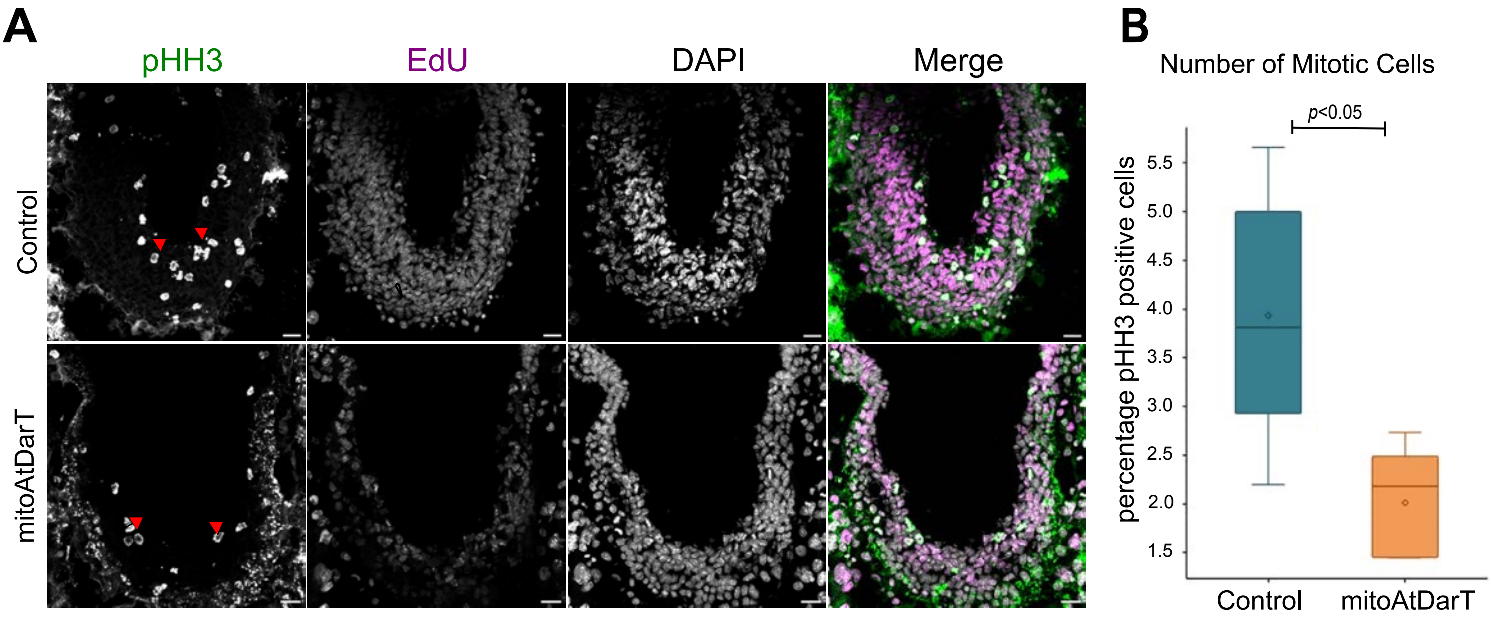
